# Osteocyte-derived erythroferrone regulates liver hepcidin during stress erythropoiesis

**DOI:** 10.1101/2024.09.27.615409

**Authors:** Vamsee D Myneni, Abhinav Parashar, Ildikó Szalayova, Eva Mezey

**Affiliations:** Adult Stem Cell Section, Craniofacial and Skeletal Diseases Branch, National Institute of Dental and Craniofacial Research (NIDCR), National Institutes of Health (NIH), Bethesda, MD 20892, USA

**Keywords:** Erythropoiesis, Osteocytes, Hepcidin, Erythroferrone, Erythropoietin receptor

## Abstract

Our knowledge of which bone marrow cells affect red cell production is still incomplete. To explore the role of osteocytes in the process we performed bulk RNAseq of osteocytes isolated from control and phlebotomized mice. The top-upregulated gene following phlebotomy was *Fam132b*, erythroferrone (*Erfe*). *Erfe* expression in osteocytes was also upregulated after erythropoietin (EPO) treatment and hypoxia *in vitro*. To explore if osteocytes contribute to the systemic ERFE levels, we generated two mouse models. We first transplanted wild-type BM in *Erfe-/-* mice creating a model where ERFE is produced in the BM but not by osteocytes. After phlebotomy, liver hepcidin suppression was significantly lower in mice where the osteocytes could not produce ERFE. To confirm that osteocytes are responsible for this difference, we generated mice lacking EPO receptors in osteocytes by crossing *Epor*^*flox/flox*^ and *Dmp1*-Cre mice. After phlebotomy, these mice showed reduced hepcidin suppression in the liver and higher circulating serum hepcidin levels compared to controls. Our work identified a novel function of osteocytes in suppressing systemic hepcidin levels during stress erythropoiesis.

## Introduction

Erythropoiesis results from a fine-tuned differentiation of hematopoietic stem cells (HSCs) into mature red blood cells (RBCs). Erythropoietin (EPO) is a master regulator of erythropoiesis, which is made by interstitial fibroblasts in the kidney (Palis, 2014). Red blood cell production requires a large amount of circulating iron. Hepcidin (Hamp), a hormone produced by the liver, regulates systemic iron levels. Down-regulation of hepcidin is required to increase the bioavailability of iron for red blood cell formation. Erythroferrone (ERFE) suppresses the production of hepcidin in the liver. Erythroferrone is produced in the bone marrow (BM) by erythroblasts in response to erythropoietic stimuli, such as blood loss, hemolysis, hypoxia, or an increase in EPO levels (Ganz and Nemeth, 2012, Ganz, 2019). The process of red blood cell development occurs along central macrophages in erythroblastic islands within the bone marrow (BM) (An and Mohandas, 2011), first described by Bessis in 1958 (Bessis, 1958).

In addition to the central macrophages, other cells such as bone marrow stromal cells (Comazzetto et al., 2019), osteoblasts (bone-forming cells) (Rankin et al., 2012), and osteoclasts (bone-resorbing cells) (Singbrant et al., 2011, Yahara et al., 2022) also support erythropoiesis either directly or indirectly (Castro-Mollo et al., 2021). Osteoblasts were found to produce sufficient EPO in the BM microenvironment to protect mice from stress-induced anemia (Rankin et al., 2012). Osteoclasts are required for the EPO-induced erythropoietic response; inhibition of osteoclast activity using bisphosphonate blunts this response (Singbrant et al., 2011). However, it’s not known if osteocytes, the most abundant bone cells (95%) that live for years unlike short-lived osteoblasts and osteoclasts (Robling and Bonewald, 2020) have a role in erythropoiesis. Osteocytes are terminally differentiated osteoblasts that reside in the bone matrix (Franz-Odendaal et al., 2006, Bonewald, 2011). Osteocytes play a vital role in skeletal homeostasis, HSC/progenitor regulation (Asada et al., 2013), myelopoiesis (Fulzele et al., 2013) and neutrophil development (Xiao et al., 2021). They secrete endocrine signals and regulate systemic phosphate homeostasis (Razzaque, 2009), adipogenesis (Fulzele et al., 2017), development of B cells and stromal cells in the thymus (Asada and Katayama, 2023).

In this work, we explored the role of osteocytes in erythropoiesis. We show that osteocytes secrete ERFE in response to erythropoietic stimuli such as phlebotomy, EPO, and hypoxia. Mice with specific deletion of EPO receptors in osteocytes showed significantly blunted hepcidin suppression in the liver and higher circulating hepcidin levels following phlebotomy. Our findings suggest that osteocytes significantly contribute to the circulating levels of ERFE to suppress hepcidin during stress erythropoiesis.

## Results

### Osteocytes produce ERFE in stress erythropoiesis

To learn about the osteocyte response to erythroid stress, 4-month-old C57BL/6 male mice were phlebotomized. After 16h, osteocyte-enriched bone tubes (referred to as “osteocytes” below) from control and phlebotomized mice were collected for RNASeq. Differentially expressed genes (DEG) were identified using START. A log^10^ change of 1 or more was used as a cutoff, with a P value of 0.05. A total of 28 DEGs were identified in osteocytes from control vs. phlebotomized mice (**Figure 1A and** see GEO dataset GSE241575). The second most significantly upregulated gene in osteocytes following phlebotomy was erythroferrone (*Erfe*) (**Figure 1A and 1B**). Since ERFE production is said to be induced by EPO (Kautz et al., 2014) we looked at the expression of the Epo receptor (*EpoR*) in osteocytes and found no difference between control and phlebotomized animals (**Figure 1C**). We validated the *Erfe* expression levels using qPCR of the same samples submitted for RNASeq. *Erfe* was upregulated 16-fold in osteocytes from phlebotomized vs control mice (**Figure 1D**). Osteocyte marker genes: dentin matrix acidic phosphoprotein (*Dmp1*), Fibroblast growth factor 23 (*Fgf23*), Matrix Extracellular Phosphoglycoprotein (*Mepe*), neuropeptide Y (*Npy*), podoplanin (*Pdpn*), Phosphate Regulating Endopeptidase Homolog X-Linked (*Phex*), and Sclerostin (*Sost*), and were not significantly different between the groups (**Figure EV1**).

**Figure 1.**
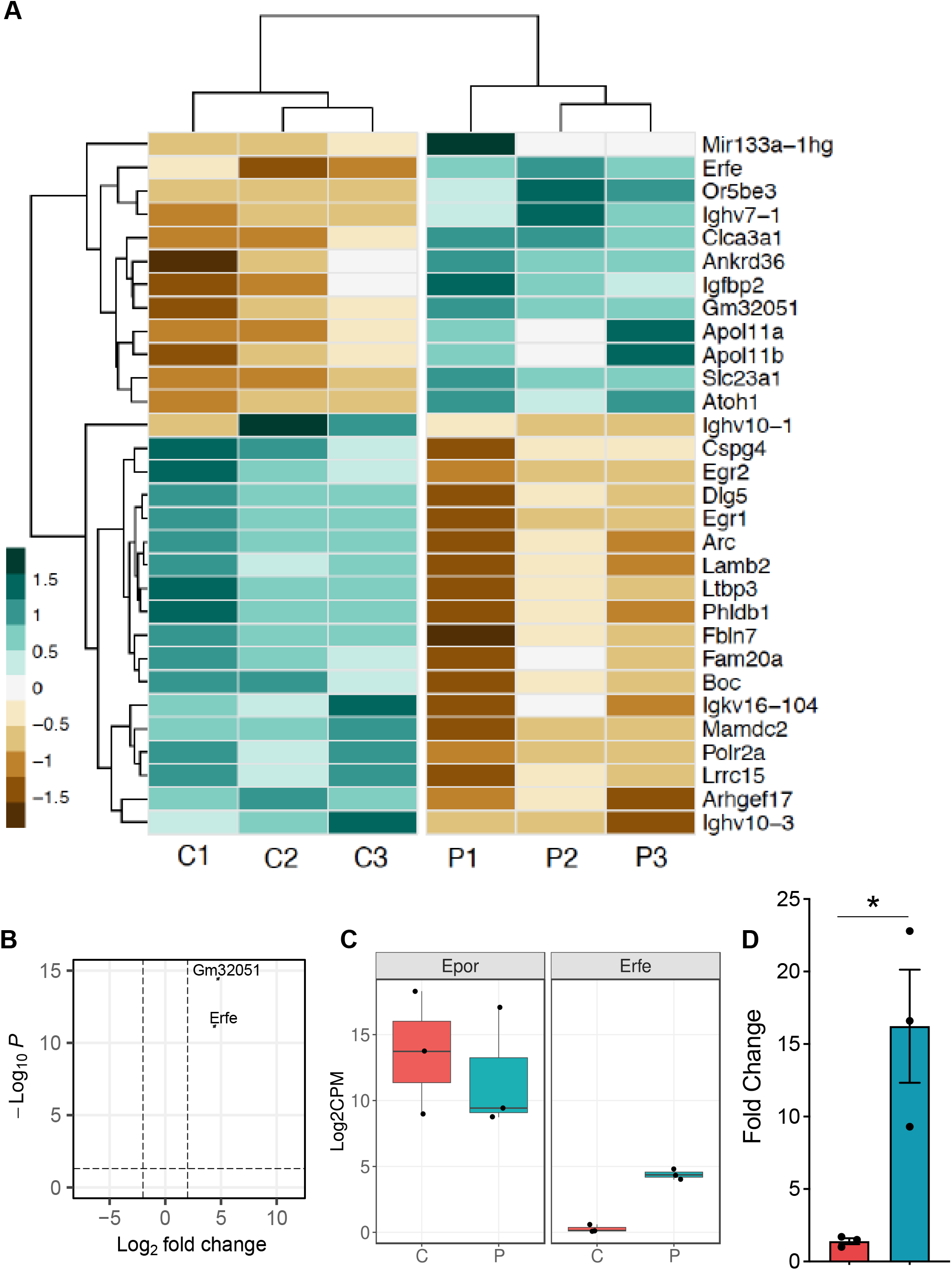
*Erfe* is the top upregulated gene in osteocytes after phlebotomy. **(A)** Heatmap showing the top 30 differentially expressed genes between control (C) and phlebotomized (P) mice of osteocytes. **(B)** The volcano plot shows that the top significant and highly expressed gene is erythroferrone (*Erfe*) after phlebotomy. **(C)** Boxplots of *Epo* receptor expression in the osteocytes are unchanged after phlebotomy, while *Erfe*’s expression is significantly increased. **(D)** qPCR confirmation of expression of *Erfe* of the RNA Seq samples (n=3, per group).

EPO has been reported to induce *Erfe* expression only in bone marrow and spleen (Kautz et al., 2014). We compared the *Erfe* expression 16h after EPO injection in total bone marrow cells, Ter119^+^ and Ter119^-^ cells, and osteocytes. In the control group, *Erfe* expression was similar in total bone marrow, Ter119^+^ cells, and osteocytes. Ter119^-^ cells had a 6-fold lower expression of *Erfe* than the other cell population. Following EPO treatment, *Erfe* expression was significantly upregulated in all four-cell populations, with highest expression in Ter119^+^ cells (6.5-fold), followed by osteocytes (4.8-fold) (**Figure 2A**).

**Figure 2.**
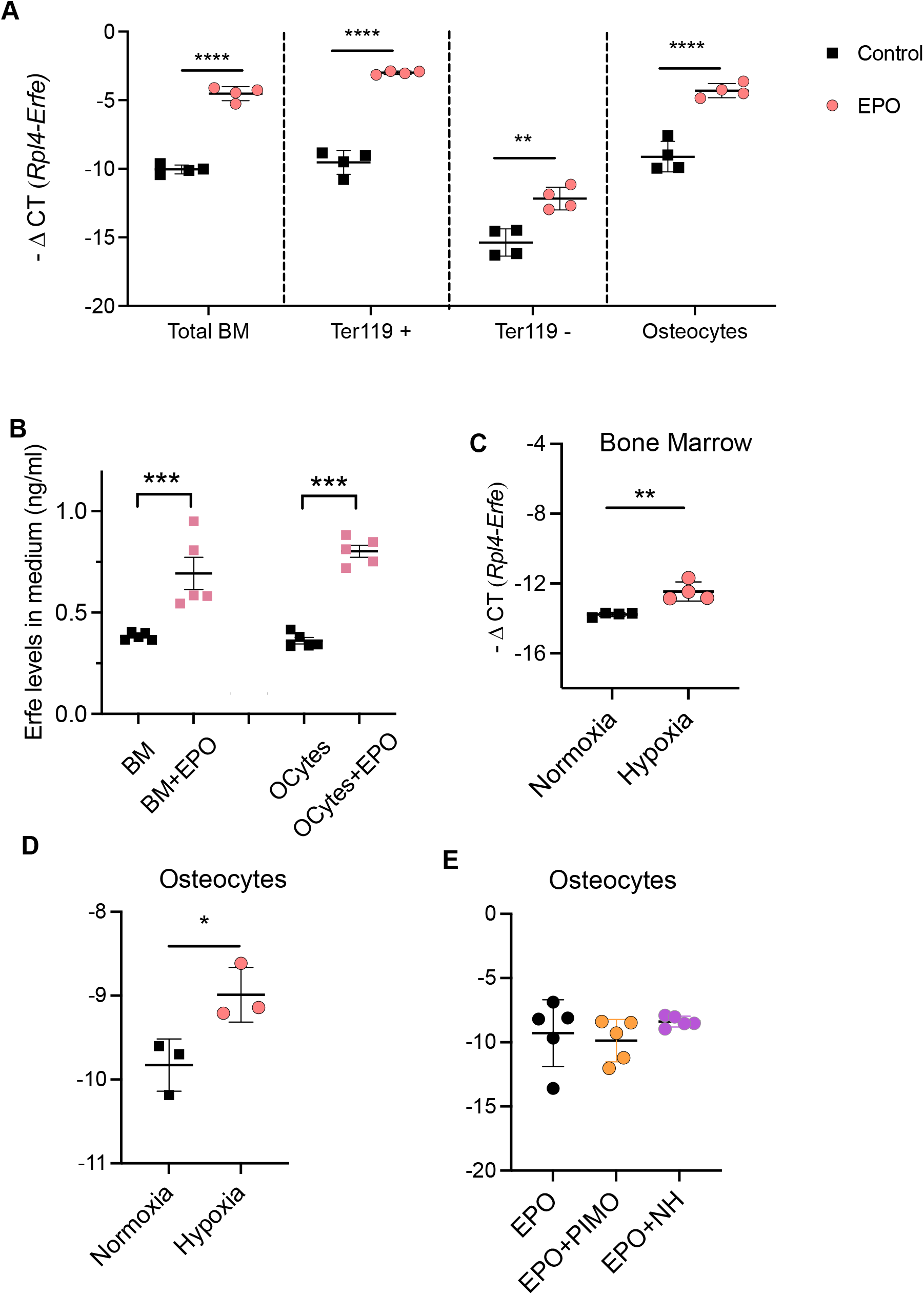
*Erfe* expression is increased in osteocytes by EPO and hypoxia. **(A)** Expression of *Erfe* in total bone marrow cells, Ter119^+^ and Ter119^-^ cells, and osteocytes, 16h after EPO injection (n=4, per group). **(B)** ERFE levels in culture media after culturing BM or osteocytes with and without EPO (10U/ml) (n=5, per group). **(C)** *Erfe* expression of bone marrow cells, cultured for 16-18h in normoxia or hypoxia (n=4, per group). **(D)** *Erfe* expression in osteocytes cultured for 16-18h in normoxia or hypoxia; (n=3). **(E)** *Erfe* expression in osteocytes following EPO stimulation and using STAT5 inhibitors pimozide (PIMO) and *N*′-((4-oxo-4H-chromen-3-yl) methylene)) nicotine-hydrazide (NH) using *in vitro* culture (n=5, per group). Data are presented as mean ± SE \**p* < 0.05, \*\**p* < 0.01, \*\*\**p* < 0.001, \*\*\*\**p* < 0.0001.

To see if osteocytes secrete ERFE in response to EPO stimulation, we cultured bone marrow cells and osteocytes with and without EPO for 3 days and analyzed the medium for ERFE using ELISA. EPO stimulation significantly increased ERFE secretion by both BM cells and osteocytes showing similar concentrations in the two populations both before and after stimulation (**Figure 2B**).

Hypoxia promotes erythropoiesis in the BM. This effect is mediated by oxygen sensors regulating the induction of EPO in interstitial fibroblasts in the kidney through hypoxia-inducible factor (HIF) (Urrutia et al., 2016, Haase, 2013). Dimethyloxalylglycine (DMOG), an inhibitor of the oxygen-sensing prolyl hydroxylases, a hypoxia mimetic agent did not induce *Erfe* expression in the bone marrow cells (Kautz et al., 2014). In order to see if hypoxia can regulate *Erfe* expression in osteocytes they were cultured in a hypoxic incubator (1.2% oxygen) alongside BM cells for 16-18h. Hypoxia significantly upregulated *Erfe* expression in both total bone marrow (**Figure 2C**) and osteocytes (**Figure 2D**). As a positive control for hypoxia culture *Vegfα* expression was analyzed in BM cells (**Figure EV 2**). To determine if osteocyte *Erfe* expression is regulated through STAT5 signaling as it is in erythroblasts, we treated osteocytes with two unrelated STAT5 inhibitors (pimozine and nicotinohydrazide labeled PIMO and NH in **Figure 2E**) for two hours before incubation with EPO for 16h. Inhibition of the JAK2-STAT5 signaling did not affect EPO-induced *Erfe* expression in osteocytes (**Figure 2E**). Overall, these results show that *Erfe* in osteocytes is regulated by both EPO and hypoxia, and *Erfe* expression in osteocytes does not seem to be regulated by the STAT5 signaling pathway.

### Osteocytes contribute to systemic ERFE levels

To see if loss of *Erfe* affects specific osteocyte marker genes, we analyzed their expression in *Erfe-/-* and *Erfe+/+* mice in steady state and following phlebotomy. In steady-state, *Dmp1* expression was significantly downregulated. However, *Fgf23* expression was significantly upregulated, and no significant changes were observed in the expression of *Mepe, Sost, Phex, E11*, and *Npy* in *Erfe-/-* compared to *Erfe+/+* mice. After phlebotomy, *Dmp1, Mepe*, and *Phex* expression were significantly downregulated, while expression of *Sost, Fgf23, E11*, and *Npy* were similar in *Erfe-/-* and *Erfe+/+* mice (**Figure EV 3**). These results suggest that *Erfe* might also play a role in skeletal mineralization; this function needs further investigation.

To see if osteocytes contribute to circulating levels of ERFE in stress erythropoiesis, we generated two mouse models. In one model, we performed bone marrow transplantation of wild-type CD45.1 donor mice to CD45.2 recipient *Erfe-/-* and *Erfe+/+* mice. The transplanted bone marrow can produce ERFE in both *Erfe-/-* and *Erfe+/+* mice, but the osteocytes in the *Erfe-/-* mice cannot. If the osteocyte-derived ERFE contributes to the suppression of liver hepcidin, we would see an increase (reduced suppression) of liver hepcidin in *Erfe-/-* mice. Three months after transplantation, the mice were subjected to phlebotomy, and liver hepcidin expression was analyzed. Suppression of hepcidin was significantly inhibited in *Erfe-/-* mice transplanted with wild-type bone marrow (**Figure 3A**) suggesting that osteocytes secrete significant levels of ERFE to suppress hepcidin during stress erythropoiesis. For the second model, we generated mice in which the EPO receptors (*Epor*) were conditionally ablated in osteocytes using the osteocyte-specific *Dmp1 cre* mice (from here on called Tg). We confirmed that *Epor* expression in osteocytes was significantly decreased in Tg compared to control mice, but not in other tissues such as spleen, liver, and bone marrow (**Figure EV 4**). Control and Tg mice were subjected to phlebotomy and hepcidin in the liver and serum were analyzed. Suppression of hepcidin expression in the liver was significantly inhibited in the Tg group compared to the control group (**Figure 3B**).

**Figure 3.**
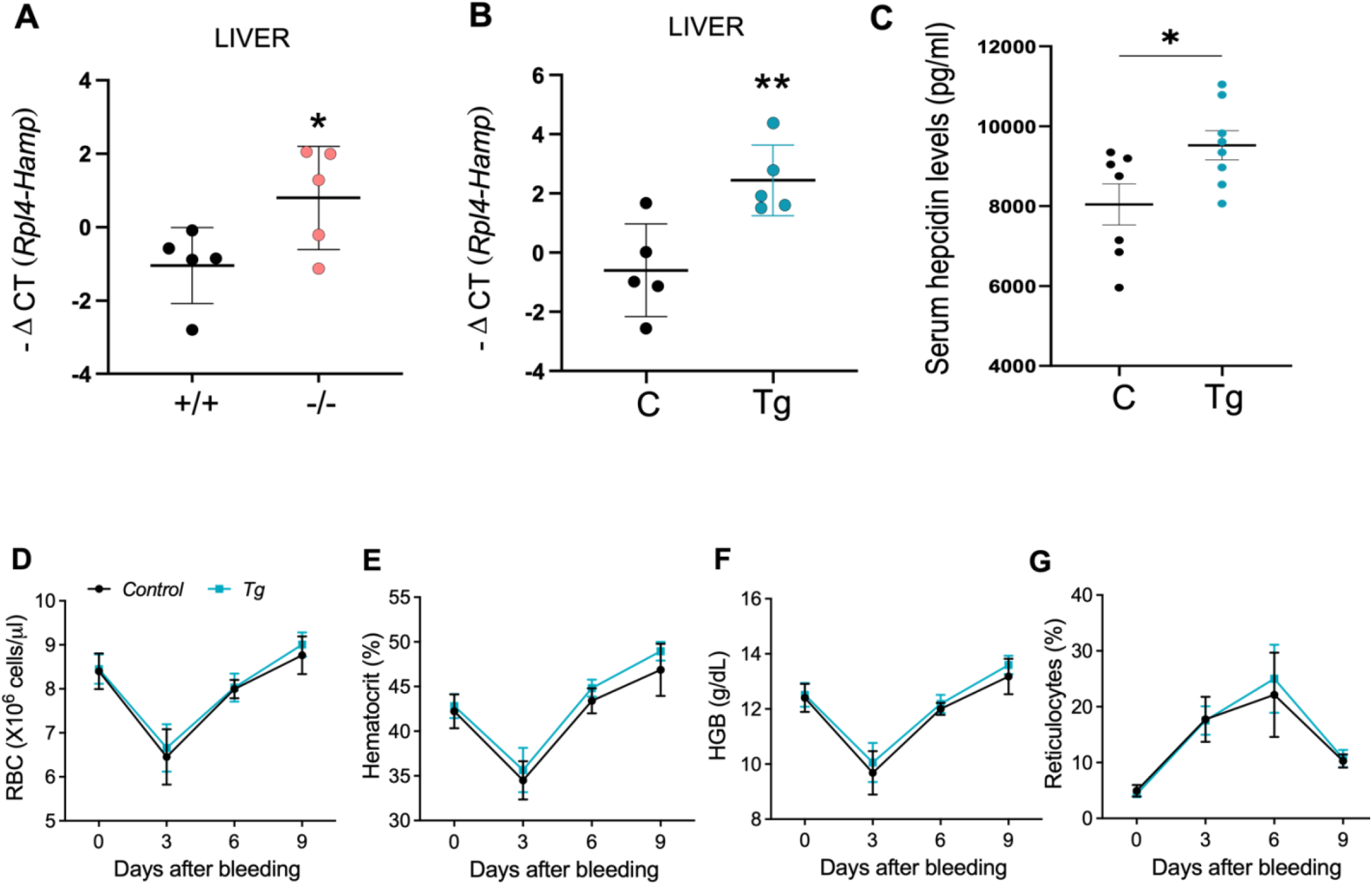
ERFE secreted by osteocytes inhibits liver hepcidin expression after phlebotomy. **(A)** *Hamp* expression in the liver after phlebotomy in *Erfe-/-* and *Erfe+/+* mice (CD45.2) transplanted with wild-type (CD45.1) mouse bone marrow; n=5, per group. **(B)** Expression of *Hamp* in the liver after phlebotomy of control and osteocyte *Epor*-deficient mice (Tg);n=5 per group. **(C)** Serum hepcidin levels after phlebotomy of control and osteocyte *Epor*-deficient mice (Tg);n=7 per group. **(D-G)** Control and Tg were subjected to phlebotomy and hematological parameters were monitored for 9 days (every 3 days), **(D)** RBC. **(E)** hemoglobin, **(F)** hematocrit, and **(G)** reticulocytes. Data are presented as mean ± SD \**p* < 0.05, \*\**p* < 0.01.

Similarly, serum hepcidin levels are higher in the Tg group compared to the control group (**Figure 3C**). To see if the loss of EPO signaling in osteocytes affects recovery after hemorrhage, control, and Tg mice were subjected to phlebotomy and monitored for 9 days. Throughout this period, there were no significant changes in RBC (**Figure 3D**), hemoglobin (**Figure 3E**), hematocrit (**Figure 3F**), and reticulocytes (**Figure 3G**). Overall, these results suggest that during stress erythropoiesis ERFE produced by osteocytes contributes to the suppression of hepcidin in the liver and that EPO receptors on osteocytes might not have a role in the recovery after hemorrhage.

## Discussion

In this work, we explored the role of osteocytes in erythropoiesis. We report a novel function of osteocytes in regulating systemic hepcidin levels through ERFE secretion during stress erythropoiesis. *Kautz et al*. (Kautz et al., 2014) reported that *Erfe* is expressed in different tissues. However, *Erfe* expression was affected only in the BM and spleen after EPO stimulation. Erythroblasts in the BM are the source of *Erfe* during stress erythropoiesis. In their study, the authors did not analyze *Erfe* expression in the bones (Kautz et al., 2014). Our work shows that *Erfe* expression in osteocytes was also stimulated by EPO and phlebotomy. This suggests that osteocytes respond to erythropoietic stimuli similarly to erythroblasts. Interestingly, *Erfe* was also expressed by other bone cells such as osteoblasts and osteoclasts (Castro-Mollo et al., 2021). However, stimulation with exogenous EPO did not upregulate the expression of *Erfe* in these cells. This observation suggests that osteocytes are the only bone cells that could regulate systemic iron levels through ERFE secretion to suppress liver hepcidin and help mobilize iron.

To study osteocytes, we first generated OEBTs (osteocytes) from mouse tibia and femur. In this culture model, the osteocytes are in their native bone matrix and retain their extensive canalicular network, which is important to maintain the phenotype and function of osteocytes *ex vivo*. Using RNASeq analysis, we identified *Erfe* as the second most highly expressed gene in osteocytes after phlebotomy. This increased expression was like that of erythroblasts reported by *Kautz et al*. (Kautz et al., 2014). To confirm this, we compared the expression of *Erfe* in erythroblasts, non-erythroblasts, and osteocytes after EPO stimulation. *Erfe* expression was increased 6.5-fold in erythroblasts and 4.8-fold in osteocytes vs. controls. Interestingly, non-erythroblasts in the BM (Ter119^-^ cells) also produce ERFE in response to EPO but much less than erythroblasts. Our work identified non-erythroblasts, and bone responds to erythropoietic stimuli like erythroblasts with different intensities.

In addition to EPO, hypoxia also stimulates erythropoiesis. Hypoxia stimulates EPO receptor expression and regulates iron metabolism (Haase, 2013). *Kautz et al*. (Kautz et al., 2014) reported that treatment of BM cells *ex vivo* using dimethyloxalylglycine (DMOG), a hypoxia mimetic agent, did not affect *Erfe* expression even though it strongly induced *Vegfα* mRNA expression. Unlike *Kautz et al*, we found that hypoxia did induce *Erfe* expression in BM and osteocytes. Similar to *Kautz et al*. (Kautz et al., 2014) *Vegfα* mRNA was also strongly induced in our BM cell cultures. The difference between *Kautz et al*. and our results might be due to their use of a hypoxia mimetic agent rather than culturing cells in a hypoxic incubator. In fact, in a recent study, *Chen et al*. compared DMOG and culturing at low oxygen, using a pheochromocytoma cell line (PC12) and found that a reduced oxygen environment is not accurately mimicked by DMOG (Chen et al., 2022). Our results also show that hypoxia is less potent in inducing *Erfe* expression than EPO. While EPO regulates *Erfe* expression through the JAK2-STAT5 signaling pathway in erythroblasts (Kautz et al., 2014), in osteocytes this does not seem to be the case. Regulation of *Erfe* expression in osteocytes and erythroblasts should be examined in more detail. Overall, our work shows that ERFE is secreted not only by erythroblasts but also by other BM cells and that osteocytes participate in suppressing liver hepcidin production in response to acute erythropoietic stimuli.

## Materials and methods

### Animal studies

Four-month-old, male C57BL/6 mice were used for bulk RNA sequencing (RNAseq). Transgenic mice with an osteocyte-specific deletion of Epor (Tg) were generated by crossing *Dmp1*-Cre mice (Jackson Laboratories, ME, US) with *Epor*^*flox/flox*^ mice (Tsai et al., 2006) (generous gift from Hong Wu, UCLA). Mice that express only Cre were used as littermate controls. *Erfe-/-* mice (Kautz et al., 2014) were a generous gift from Tomas Ganz, UCLA. For erythropoietic stimulus, mice were phlebotomized by retro-orbital puncture (500 μl) or treated with a single dose of 200U of human EPO (NDC55513-126-10) Epogen, Amgen). Analysis was done 16 hours later. RNA was prepared from livers as described below and serum hepcidin levels were measured using ELISA according to the manufacturer’s instructions (NOVUS Ms hepcidin ELISA, cat#NBP2-82129). Recovery from anemia was monitored every 3 days after phlebotomy for 9 days. Blood (50 μl) was collected by a retro-orbital puncture at each time point. Complete blood counts were obtained using the IDEXX ProCyte Dx hematology analyzer. Animal housing and maintenance were in full compliance with NIH criteria for the care and use of laboratory animals. The Animal Care and Use Committee of the National Institute of Dental and Craniofacial Research (NIDCR), NIH, approved all procedures.

### Osteocyte enriched bone tube (OEBT) isolation and culture

OEBTs were generated from the tibia and femur. The bone marrow was isolated by centrifugation as previously described (Myneni et al., 2021). Osteocytes were isolated by a modified protocol of Stern and Bonewald (Stern and Bonewald, 2015). The bones were subjected to sequential digestion using collagenase buffer (αMEM media, HEPES (25mM), BSA (0.1%), Penicillin-streptomycin (1%), Collagenase type IV (2 mg/ml), Dnase-40 ug/ml) and EDTA buffer (PBS, BSA (0.1%), EDTA 5mM) as follows. Collagenase, 20 min; EDTA, 20 min; collagenase, 1h; collagenase, 1h; EDTA, 45 min. After the final digestion, the tubes were washed and used for RNA extraction or cell culture. For RNA expression analysis, the tubes were cultured in αMEM medium, 0.2% polyvinyl alcohol (PVA), and Penicillin-streptomycin (1%) and stimulated with EPO (10 U/ml). For ERFE culture medium analysis, total BM cells are cultured in IMDM with 10% FBS. OEBTs are cultured in αMEM medium with 10% FBS for three days and ERFE was measured by sandwich Elisa (Biomatik, Cat#EKF58568).

For STAT5 signal inhibition, the OEBTs were pretreated for 2 hours with two different STAT5 inhibitors - pimozide (10μM-Sigma; Cat#P1793), or *N*′-((4-oxo-4H-chromen-3-yl) methylene) nicotinohydrazide (200μM-Cayman chemicals; Cat#15784) before EPO stimulation. After 16h the tubes were collected to study RNA expression as described above.

### Isolation of bone marrow red blood cells

Bone marrow cells were separated into Ter119^+^ and Ter119^−^populations using magnetic beads. Total bone marrow cells (1×10^7^) were stained with anti-mouse Biotin-Ter119 antibody (Thermo Fisher Scientific #13-0114-82) and MagCellect streptavidin ferrofluid (R & D; MAG999B) magnetic beads were used for selection. Ter119^+^ cells were collected using a MagCellect magnet; the flow-through contained Ter119^−^cells. All the cells underwent RNA analysis.

### Bone marrow transplantation

For bone marrow transplantation 1×10^6^ bone marrow cells were isolated from CD45.1-positive WT mice given intravenously to lethally irradiated (10Gy) CD45.2 *Erfe+/+* and *Erfe-/-* recipient mice at 6–8 weeks of age. Transplanted mice were given sulfa-methoxazole- and trimethoprim-containing (TMS) water for 3 weeks, followed by regular water. Bone marrow engraftment was confirmed by flow cytometric analysis of CD45.2-positive donor-derived cells in the peripheral blood of recipient mice. Three months after bone marrow transplantation the mice were subjected to phlebotomy. Sixteen hours later the livers of mice were collected and analyzed for hepcidin expression.

### RNA extraction and qPCR

RNA extraction from bone tubes, liver, and cells was done using TRIzol LS (Invitrogen) and Direct-zol RNA purification kits (Zymo, CA, USA) per the manufacturers’ instructions. Bone tubes were homogenized using Lysis Matrix (MP Biomedicals, USA). The cDNA synthesis was performed with a High-Capacity cDNA reverse transcription kit (Applied Biosystems). QPCR was done with GreenLink High-ROX QPCR master mix (BioLink); the primers used are osteocyte marker genes (Asada et al., 2013), *Hamp, Erfe, Rpl4, Vegfα* (Kautz et al., 2014), *Epor (Suresh et al*., *2020)*. The reactions were run on a QuantStudio 5 real-time PCR system (Applied Biosystems). mRNA transcript abundance was normalized to that of *Tub1a* or *Rpl4* reference (housekeeping) genes. Gene expression (−ΔC_T_) is the threshold cycle (C_T_) for the gene of interest minus the C_T_ of the reference gene.

### Bulk RNA Sequencing

RNA was converted into cDNA using the SMART-seq UltraLow v4 kit (Clontech, Mountain View, CA). Illumina cDNA sequencing libraries were prepared using the Nextera XT kit (Illumina, San Diego, CA). Libraries were sequenced on an IlluminaHiSeq 1500 sequencer, in the 126bp paired-end mode. Raw sequences underwent initial QC analysis and were then aligned to the GRCm38 version of the mouse genome with STAR v2.5.2a. Raw gene read counts produced by STAR were filtered to remove low-expressing genes and further processed in R (R Development Core Team, 2011) using the EdgeR package (McCarthy et al., 2012) and the START application (Nelson et al., 2017).

## Statistical Analysis

Statistical analyses were performed with Prism 8 (GraphPad Software, Inc.) using unpaired t-test. Significance is indicated by ∗p < 0.05, ∗∗p < 0.01, ∗∗∗p < 0.001, ∗∗∗∗p < 0.0001.

## Data Sharing Statement

The RNAseq data are available under GEO accession GSE241575.

## Acknowledgments

The above study was supported by the intramural research program (IRP) of NIDCR, NIH under project ZIA DE000714. The authors acknowledge the help of Daniel Martin, PhD for analyzing the genetic data set and the employees of the NIDCR veterinary core (VRC) for their help with taking care of the animals (Genomics and Computational Biology Core: ZIC DC000086 and Veterinary Resources Core: ZIC DE000740-05). All animal procedures have been approved by NIDCR’s Animal Care and Use committee under ASP 19-1000.

## Authorship Contributions

VDM designed the study, performed experiments, performed data analysis, wrote the draft, prepared draft figures, AP performed in vitro experiments, helped analyze data and participated in finalizing the text and figures; ISz performed irradiation and BM transplants, was responsible for breeding and genotyping mice and performed all ELISA experiments, EM oversaw design, analyzed data, and performed more detailed RNAseq analysis (with Daniel Martin), prepared final figures, and wrote the final version of the paper.

## Disclosure of Conflicts of Interest

The authors have no conflict of interest.

## Expanded View Figure legends

Expendable View Figure 1. RNASeq analysis of osteocytes after phlebotomy

Osteocyte specific gene expression based on bulk RNAseq results comparing control to phlebotomized mice.

Expendable View Figure 2. qPCR expression of *vegfα* in bone marrow cells in normoxia and hypoxia confirming the hypoxic effect of VEGFa (positive control).

Expendable View Figure 3. Osteocyte marker gene of WT and *Erfe-/-* mice. (A) Steady-state osteocyte specific marker genes (B) Osteocyte specific marker genes after phlebotomy.

Expendable View Figure 4. *Epor* in different tissues of wild type and Tg mice confirming the specific deletion in osteocytes.

